# Valence-specific ensembles in the laterodorsal tegmentum encode salient stimuli and modulate motivated behavior

**DOI:** 10.64898/2026.01.28.702313

**Authors:** E Teixeira, R Bastos-Gonçalves, LAA Aguiar, L Royon, AC Faria, R Jungmann, D Vilasboas-Campos, M Cunha-Macedo, AV Domingues, C Soares-Cunha, R Correia, SP Fernandez, J Barik, AJ Rodrigues, B Coimbra

## Abstract

The laterodorsal tegmentum (LDT) is a brainstem hub that integrates sensory and motivational signals to regulate adaptive behavior. While LDT neurons are known to modulate reward and aversion, whether salient stimuli recruit distinct neuronal ensembles within this structure remains unknown. Here, combined cell-type-specific calcium imaging, activity-dependent genetic tagging (TRAP2), and optogenetic reactivation to investigate how rewarding and aversive stimuli recruit and functionally define LDT neurons. Notably, single exposures to cocaine or shock in TRAP2;Ai14 mice labeled spatially and neurochemically distinct ensembles, with minimal overlap.

We used fiber photometry and TRAP2 system to tag active neuronal ensembles in the LDT during cocaine (coca-LDT) and foot shock (shock-LDT) exposure, expressing GCaMP8m (green) in these neurons and, simultaneously, sRGECO (red) in the whole LDT. Our results demonstrate coca-LDT activation by physical aversive events and valenced odours, with reduced activity in response to rewarding liquids. Shock-LDT showed activation by physical aversive events, odours, and shock-predictive cues. Additionally, optogenetic reactivation of cocaine-TRAPed ensembles in a two-choice operant task biased action selection toward stimulation-paired responses, whereas shock-TRAPed ensemble activation did not drive avoidance. These findings identify functionally segregated LDT ensembles recruited by opposing motivational stimuli and reveal a causal role for reward-activated brainstem ensembles in shaping behavior. This functional segregation may contribute to the brain’s ability to differentiate stimulus types and, to some extent, valence experiences. Our results may provide evidence on how the LDT influences decision-making processes in addiction and anxiety disorders, potentially paving the way for novel therapeutic approaches.

## Introduction

Emotionally salient experiences, whether aversive or appetitive, drive behavioral adaptation by recruiting discrete neural circuits that assign value and guide action. These processes rely on distributed brain networks, including cortical, limbic, and midbrain areas, which encode motivationally relevant stimuli and orchestrate flexible responses to environmental demands(refs). While much is known about how forebrain structures such as the amygdala, prefrontal cortex, and nucleus accumbens (NAc) represent valence, the contribution of brainstem structures, notably the laterodorsal tegmentum (LDT), remains poorly understood.

The LDT is a pontine nucleus composed of cholinergic, glutamatergic, and GABAergic neurons that project broadly to forebrain regions implicated in reward, aversion, arousal, and cognitive function, including the ventral tegmental area (VTA), thalamus, hypothalamus, and NAc (refs). Its anatomical position and connectivity suggest a critical integrative role in shaping behavior based on environmental and internal states. Indeed, accumulating evidence implicates LDT neurons in both reward and aversion processing. Cholinergic and glutamatergic neurons in the LDT have been shown to drive reinforcement learning and preference-related behavior ^1–5^, whereas GABAergic neurons exert more heterogeneous effects, including aversive responses^6,7^. For example, optogenetic activation of LDT cholinergic projections to the VTA or NAc promotes place preference and reward-seeking behavior, whereas inhibition has the opposite effect. This modulation has been shown to impact conditioned place preference for cocaine, a drug of abuse^8^. Stimulation of GABAergic projections can induce aversion or suppress drug intake^2,3,5,9,10^. At the circuit level, the LDT receives input from both reward-related regions (e.g., prefrontal cortex, lateral hypothalamus) and aversion-related regions (e.g., lateral habenula, periaqueductal gray)^6,11,12^, supporting its role as a node for valence integration.

Despite this, it remains unknown whether specific LDT ensembles are selectively recruited by stimuli of opposing valence, and whether such ensembles are sufficient to influence behavioral output. Much of what is known about valence processing in the LDT has been inferred from population-level manipulations using cell-type-specific tools. However, distinct neuronal ensembles, descried as sparse populations of neurons activated during specific experiences, are increasingly recognized as key units of emotional memory and motivational encoding across the brain^13^.

To investigate whether the LDT exhibits ensemble-level organization during exposure to salient stimuli, we used the TRAP2 (Targeted Recombination in Active Populations 2) model to genetically tag and manipulate neurons active during either appetitive (cocaine) or aversive (footshock) conditions. We then characterized the neurotransmitter identity, functional responsiveness, and assessed their behavioral relevance through optogenetic reactivation.

Our findings reveal that rewarding and aversive stimuli recruit largely non-overlapping ensembles in the LDT. By using *in* vivo fiber photometry, we simultaneously recorded calcium activity of overall cells in the LDT and in stimulus defined populations during aversive and rewarding tasks. These results showed divergent activity patterns during stimulus exposure. Finally, optogenetic reactivation of cocaine-recruited LDT ensembles was sufficient to bias action selection in an operant two-choice task, whereas reactivation of aversion-recruited ensembles did not induce avoidance.

These results demonstrate that the LDT encodes salient motivational stimuli through valence-specific ensemble recruitment, and that reward ensembles within the LDT are causally involved in shaping motivated behavior. Our study extends the concept of functional engrams to the brainstem and highlights the LDT as an integral node for motivational salience processing.

## Results

### Genetically defined LDT populations are recruited in response to different modalities of valence stimuli

Previous studies assessed the effect of modulation of different populations (cholinergic, glutamatergic or GABAergic) in preference-related behaviours, leading to often opposing behavioral outcomes dependent on the nature of the population ^3,7,14^. To determine if exposure to opposing valence stimuli specifically alters the response dynamics of each LDT population, cre-dependent GCaMP8m was injected into the LDT of ChAT-, vGluT-, and vGAT-cre mice to selectively target each neuronal population (Figure 1A). Following recovery, animals were exposed to stimuli of distinct valence, including TMT, condensed milk, cocaine exposure and shock delivery (Figure 1B).

**Figure 1.**
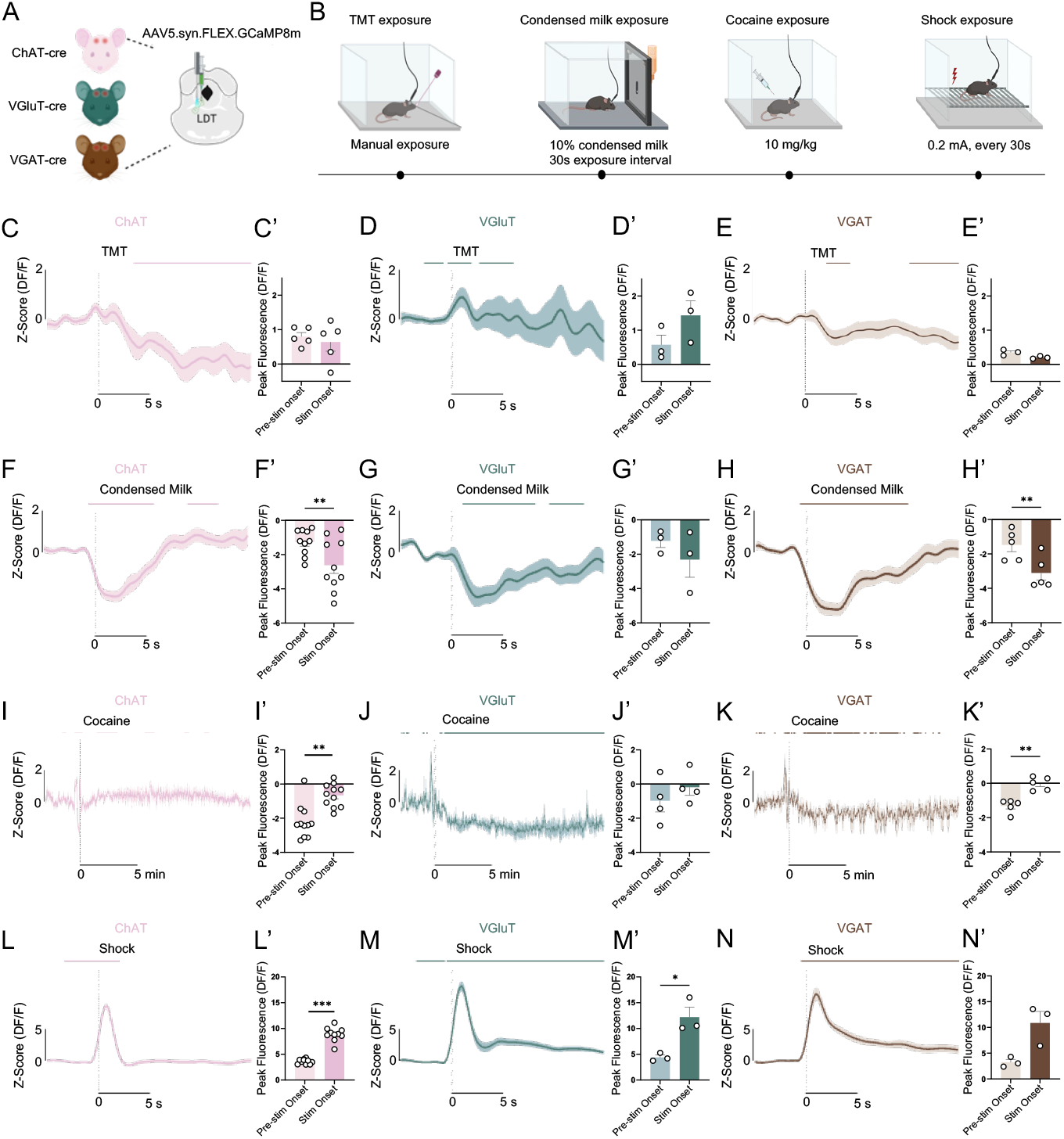
Response of different LDT populations to varied stimuli. (A) Schematic representation of photometry readings of LDT neuronal populations, cholinergic (ChAT, n = 11), glutamatergic (vGluT, n = 4) or GABAergic (vGAT, n = 5) with expression of GCaMP8m calcium sensor, respectively, in the LDT of ChAT-cre, VGluT-cre or vGAT-cre mice and optic fiber implantation. (B) Sequence of stimuli exposure. (C-E’) LDT populations response to Trimethylthiazoline (TMT – aversive odour), represented as Z-score of the photometry recordings (C, D, E) and its peak fluorescence analysis (D’, E’, F’), showing no differences in activity aligned to the odour exposition. (F-H’) LDT populations response to condensed milk consumption (rewarding liquid), represented as Z-score of the photometry recordings (F, G, H), that show an activity decrease in all populations, and the Z-score negative fluorescence analysis (F’, G’, H’), evidencing a decrease in ChAT and VGAT populations (paired Student’s t test, ChAT: t(9) = 4.138, p = 0.0025; VGAT: t(2) = 5.149, p = 0.0067). (I-K’) LDT populations response to cocaine injection (rewarding substance), represented as Z-score of the photometry recordings (I, J, K), showing little change after cocaine injection, as well as peak fluorescence analysis (I’, J’, K’), showing activity increases in ChAT and VGAT population (Wilcoxon test, ChAT: W = 66.00, p = 0.0010; paired Student’s t test, VGAT: t(4) = 5.519, p = 0.0053). (L-N’) LDT populations response to footshock (nociceptive aversive event), represented as Z-score of photometry (L, M, N), showing an activity increase in all LDT populations, and peak fluorescence analysis (L’, M’, N’), that evidence the activity analysis in ChAT and VGluT (paired Student’s t test, ChAT: t(9) = 16.61, p < 0.0001; VGluT: t(2) = 5.230, p = 0.0347) and an increase tendency in VGAT (t(2) = 4.241, p = 0.0513). Z-score data are represented as trial mean (darker line) ± SEM (lighter area). Lines over Z-score graphs indicate periods of significant difference from baseline as defined by bootstrap 95% CI. Data is presented as mean ± SEM. * p < 0.05, ** p < 0.01, *** p < 0.001.

Exposure to TMT, an innately aversive odor used to mimic predator presence, did not elicit significant changes in the activity in any of the LDT neuronal populations examined (ChAT, vGluT, or vGAT) (Figure 1C, D, E), a result further supported by the absence of significant effects in peak fluorescence analysis (Figure 1C’, D’, E’).

Condensed milk exposure was associated with an overall decrease in activity (Figure 1F, G, H), with peak fluorescence analysis revealing significant reductions in calcium activity in both ChAT and VGAT populations (Figure 1F’, G’, H’).

To test whether LDT neurons respond to pharmacologically induced reward states, we recorded calcium activity during intraperitoneal, i.p, injections of cocaine (20 mg/kg). Cocaine administration (20mg/Kg) did not produce marked changes in population-level activity of vGluT-cre animal (Figure 1J, J’). Both ChAT and vGAT populations increased activity upon cocaine exposure (Figure 1I, K), as observed by peak fluorescence analysis (Figure 1I’, K’).

Finally, foot shock induced robust increases in activity across all LDT populations, an effect that was consistently reflected in the bootstrap 95% CI and peak fluorescence analyses (Figure 1L-N’). Collectively, these data indicate that LDT neuronal subtypes are differentially recruited by salient stimuli depending on both valence and sensory modality.

### Valence representation in LDT comprises distinct ensemble populations

Considering that all LDT populations responded to either positive or negative valence stimuli, and to different modalities of stimuli, we aimed to characterize how specific neuronal ensembles would respond to stimuli with the same valence or opposite. We used the TRAP2;Ai14 mouse model^15^, with a Fos locus to drive the expression of a tamoxifen-inducible Cre-recombinase (Fos 2A-iCreERT2/TRAP2 allele), that in the presence of the 4-OHT-inducible Cre-recombinase (CreRTT2), enables permanent expression of effector fluorescent protein in recruited cells, tdTomato (Figure 2A). Mice were habituated to i.p. injections for 7 days, followed by cocaine (20 mg/Kg) injection or shock exposure (1.5mA, 1s x 10) on TRAPing day with i.p. administration of 4-OHT to induce expression of tdTomato. After TRAPing, each animal was exposed to a different stimulus to then perform c-fos quantification via immunostaining. This allows to assess the number of cells activated by two separate stimuli in the same animal. Thus, cells activated by a first stimulus in the presence of tamoxifen will be tdTomato-positive (tdTomato+) and cells activated by a second stimulus later will be endogenous c-Fos+ (green) (Figure 1B), allowing the study of overlap or segregation between ensembles representing separate events. Exposure to the same stimulus, i.e., animals exposed to shock-shock (∼ 37%) and cocaine-cocaine (∼36%) showed increased co-localization (Figure 2C). In contrast, opposing stimuli, cocaine-shock, only presented 11% of overlap, suggesting that LDT recruits distinct neuronal ensembles, depending on stimulus valence.

**Figure 2.**
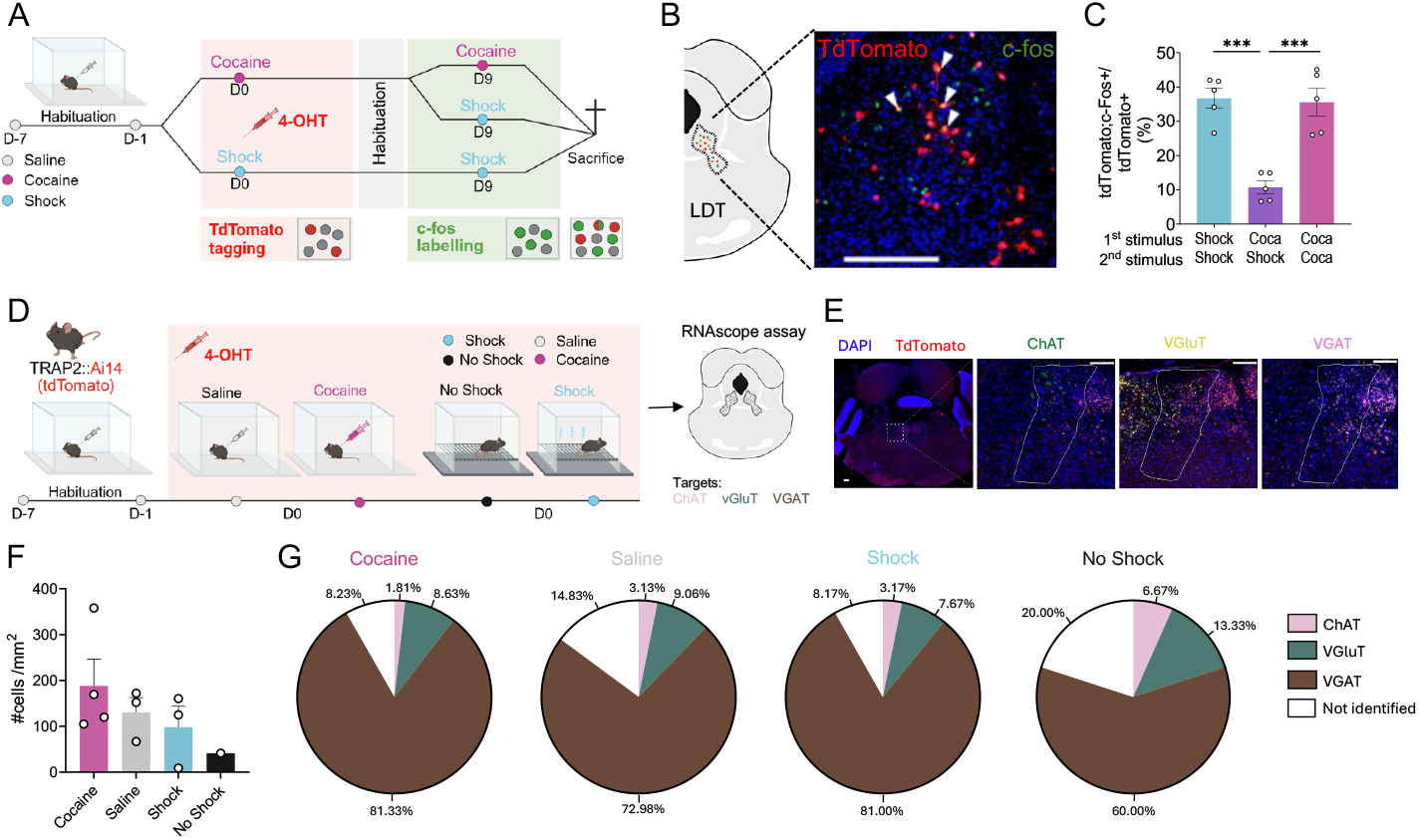
Valence specificity and neurotransmitter content of cocaine- and shock-recruied LDT ensembles. (A) Strategy of recruitment of activated cells in TRAP2;Ai4 mice (n = 15) by cocaine (n = 10) or shock (n = 5) and c-fos detection of activated cells by the same stimuli a second time. (B) Histology example of TdTomato (from model) and c-fos immunostaining in the LDT. (C) Co-localisation results of c-fos stained cells on the total TdTomato stained cells, showing higher colocalisation for exposure to stimuli of same valence (one-way ANOVA, F(2,12) = 23.07, p < 0.0001; *post hoc*: coca-shock vs shock-shock: t = 6.0009, p = 0.0002, coca-shock vs coca-coca: t = 5.747, p = 0.0003). (D) Strategy for cocaine-, saline-, shock- or no shock-TRAPping that allowed expression of TdTomato in recruited cells and RNAscope assay to identify the neurotransmitter content of TRAPped neurons. (E) Histological examples of RNAscope staining of different LDT populations (ChAT, vGluT and vGAT) and TdTomato staining of recruited cells. (F) TdTomato cell density in the LDT after exposure to different stimuli, showing no difference in recruitment cells number by different stimuli. (G) RNAscope cellular populations and stimuli-recruited cells co-localisation results, depicted as share of each LDT population based on neurotransmitter content on the total tdTomato stained cells, showing vGAT predominance in recruited cells in all populations, and higher content of VGluT recruited cells in animals exposed to cocaine or shock. Scalebars represent 200 µm. Data is presented as mean ± SEM. * p < 0.05, ** p < 0.01, *** p < 0.001.

To determine the genetic nature of coca- or shock-recruited ensembles, TRAP2;Ai14 animals were exposed to the same opposing stimuli, with the addition of control groups for each stimulus (saline injection as control for cocaine ensembles or exposure to behavior box, No shock group, as control for shock ensemble) (Figure 2D). RNAscope *in situ* hybridization data showed that cocaine- or shock-activated ensembles recruited more cells than respective controls (Figure 2F). Using probes to target cholinergic (ChAT), glutamatergic (vGlut2) and GABAergic (GAD2) neurons, the majority of cells in each ensemble were GABAergic, though all 3 populations were present for every stimulus exposure.

### Aversive stimuli recruits LDT ensembles

Having the characterisation of the nature of the LDT cocaine- and shock-activated neuronal populations, we further aimed to describe the response of these recruited populations to different valence stimuli. To this end, we injected cre-dependent GCaMP8m and generally expressed sRGECO into the LDT of TRAP2 mice and used the same TRAPping strategy as previously described, expressing GCaMP8m in cocaine- or shock-recruited and the general LDT population, respectively (Figure 3A-B).

**Figure 3.**
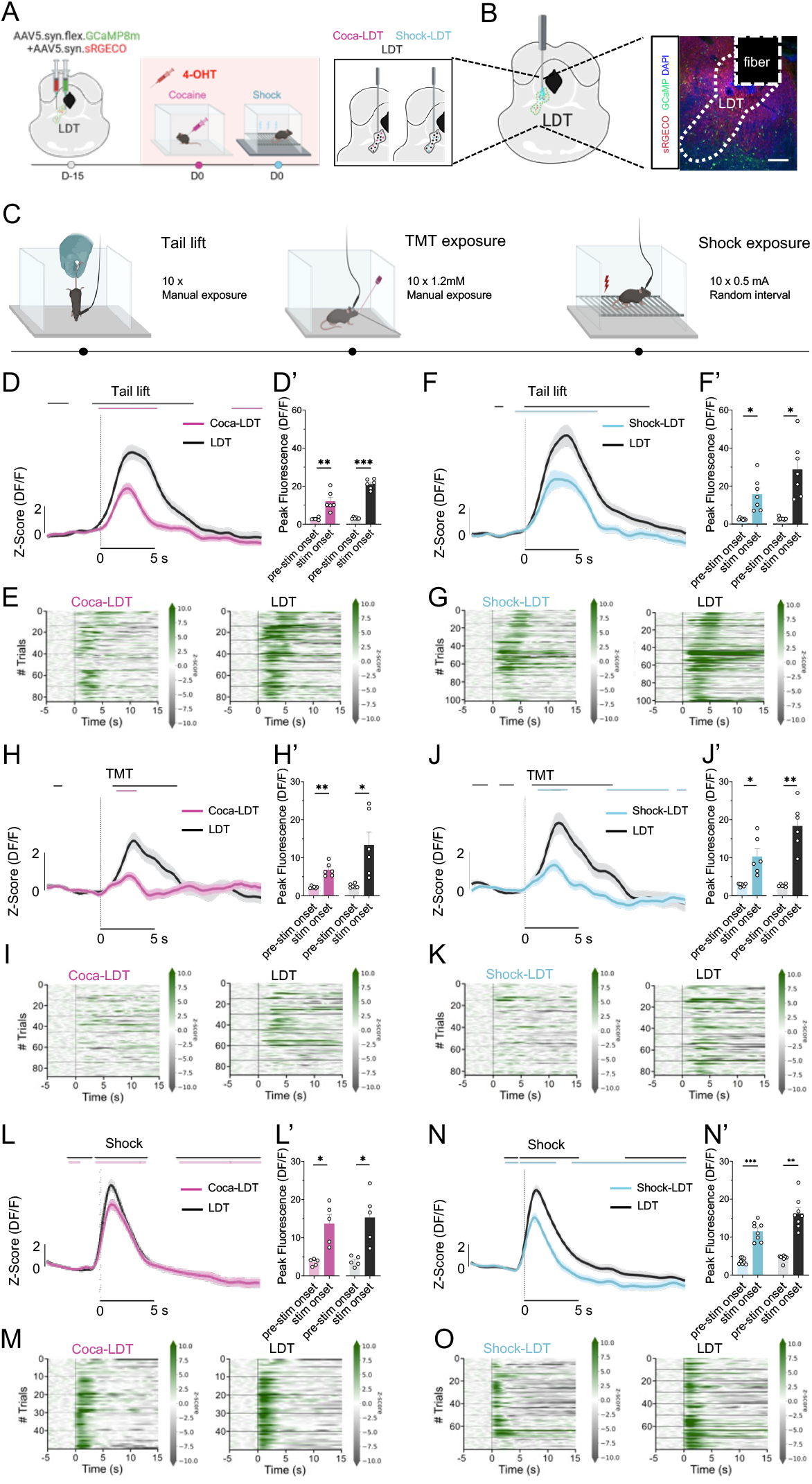
LDT and shock- and cocaine-recruited LDT neuronal populations’ response to negative stimuli. (A) Schematic representation of injection of cre-dependent GCaMP8m and general expressed sRGECO in the LDT (LDT) of TRAP2 mice (n = 20) and strategy of recruitment of neurons activated by cocaine ( n = 10; coca-LDT) or shock (n = 10, shock-LDT). (B) Histological checking of viral expression. (C) Sequence of negative stimuli mice were exposed to. (D-E) Coca-LDT and LDT response to tail lift, showing an activity increase aligned to lift moment, depicted in both z-score of the DF/F (D) and peak fluorescence analysis (D’, paired Student’s t test, coca-LDT: t(5) = 4.150, p = 0.0089, LDT: t(5) = 17.34, p < 0.0001), with trial response detail depicted in heatmap (E). (F-G) Shock-LDT and LDT response to tail lift, showing an activity increase aligned to lift moment, depicted in both z-score of the DF/F on PSTH of the stimulus (F) and peak fluorescence analysis (F’, paired Student’s t test, shock-LDT: t(6) = 4.179, p = 0.0058, Wilcoxon test, LDT: W = 28.00, p = 0.0156); trial response detail depicted in heatmap (G). (H-I) Coca-LDT and LDT response to TMT, showing an activity increase aligned to TMT exposure, depicted in both z-score of the DF/F on PSTH of the stimulus (H) and peak fluorescence analysis (H’, paired Student’s t test, coca-LDT: t(5) = 6.649, p = 0.0012, LDT: t(5) = 3.189, p = 0.0243), with trial response detail depicted in heatmap (E). (J-K) Shock-LDT and LDT response to TMT, showing an activity increase aligned to TMT exposure, depicted in both z-score of the DF/F on PSTH of the stimulus (J) and peak fluorescence analysis (J’, Wilcoxon test, shock-LDT: W = 21.00, p = 0.0312, paired Student’s t test, LDT: t(5) = 6.445, p = 0.0013); trial response detail depicted in heatmap (K). (L-M) Coca-LDT and LDT response to shock, showing an activity increase aligned to shock administration, depicted in both z-score of the DF/F on PSTH of the stimulus (L) and peak fluorescence analysis (L’, paired Student’s t test, coca-LDT: t(4) = 4.270, p = 0.0129, LDT: t(4) = 3.878, p = 0.0179), with trial response detail depicted in heatmap (M). (N-O) Shock-LDT and LDT response to shock, showing an activity increase aligned to shock administration, depicted in both z-score of the DF/F on PSTH of the stimulus (N) and peak fluorescence analysis (N’, paired Student’s t test, shock-LDT: t(7) = 12.46, p < 0.0001, Wilcoxon test, LDT: W = 36.00, p = 0.0078); trial response detail depicted in heatmap (O). Scalebars represent 200 µm. Z-score data are represented as trial mean (darker line) ± SEM (lighter area). Lines over Z-score graphs indicate periods of significant difference from baseline as defined by bootstrap 95% CI. Bar data are represented as animal mean ± SEM. * p < 0.05, ** p < 0.01, *** p < 0.001

Animals were subsequently exposed to different aversive stimuli, namely tail lift, TMT exposure and shock exposure (Figure 3C).

Across all aversive stimuli, both the general LDT population and cocaine- and shock-recruited ensembles exhibited a significant increase in the activity during stimulus presentation(Figure 3D-O). This effect was further supported by peak fluorescence analysis (Figure 3D-O).

### Appetitive stimuli recruits LDT ensembles

After determining how LDT shock- and cocaine-recruited ensembles response to aversive stimuli, we assessed their engagement by appetitive stimuli (Figure 4A).

**Figure 4.**
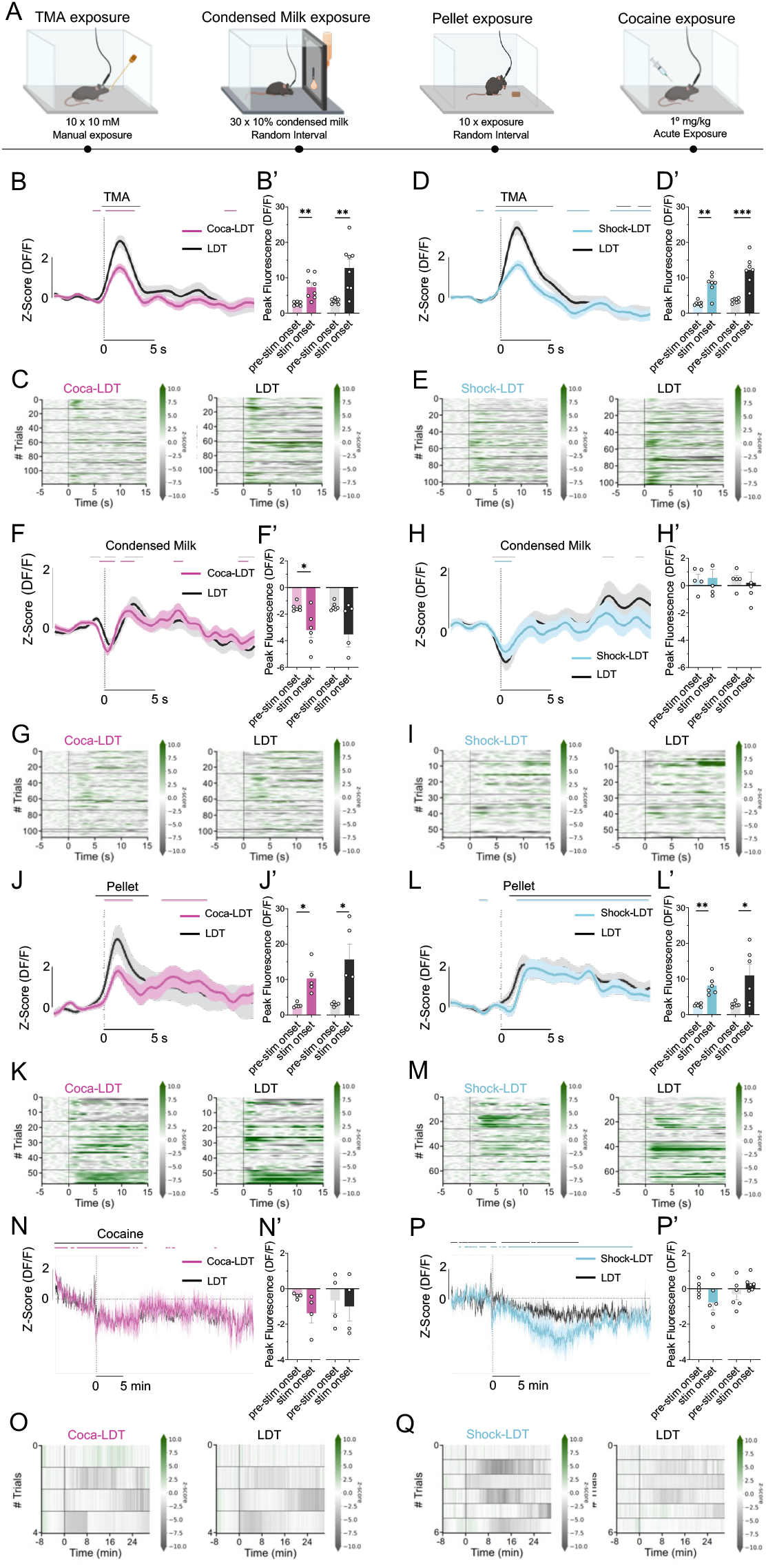
LDT and shock- and cocaine-recruited LDT neuronal populations’ response to positive stimuli. (A) Sequence of positive stimuli mice were exposure to. (B-C) Coca-LDT and LDT response to TMA, showing an activity increase aligned to TMA exposure, depicted in both z-score of the DF/F (B) and peak fluorescence analysis (B’, paired Student’s t test, coca-LDT: t(7) = 4.062, p = 0.0048, LDT: T(7) = 4.096, p = 0.0046), with trial response detail depicted in heatmap (C). (D-E) Shock-LDT and LDT response to TMA, showing an activity increase aligned to TMA exposure, depicted in both z-score of the DF/F on PSTH of the stimulus (D) and peak fluorescence analysis (D’, paired Student’s t test, coca-LDT: t(6) = 5.930, p = 0.0010, LDT: t(6) = 6.150, p = 0.0008); trial response detail depicted in heatmap (E). (F-G) Coca-LDT and LDT response to condensed milk, showing an activity decreased of the coca-LDT neurons aligned to condensed milk consumption, depicted in z-score of the DF/F on PSTH of the stimulus (F) and analysis of the negative peak fluorescence for coca-LDT, as it shows a decrease tendency in LDT (F’, paired Student’s t test, coca-LDT: t(5) = 3.555, p = 0.0163, LDT: t(5) = 2.361, p = 0.0647), with trial response detail depicted in heatmap (G). (H-I) Shock-LDT and LDT response to condensed milk, showing an activity decrease aligned with condensed milk consumption, depicted in both z-score of the DF/F on PSTH of the stimulus (H) but not on analysis of the negative peak fluorescence (H’); trial response detail depicted in heatmap (I). (J-K) Coca-LDT and LDT response to food pellets, showing an activity increase aligned to the food pellet consumption, depicted in both z-score of the DF/F on PSTH of the stimulus (J) and peak fluorescence analysis (J’, Wilcoxon, coca-LDT: W = 15.00, p = 0.0625, paired Student’s t test, LDT: t(4) =3.050, p = 0.0380), with trial response detail depicted in heatmap (K). (L-M) Shock-LDT and LDT response to food pellets, showing activity increase to food pellet consumption, depicted in z-score of the DF/F on PSTH of the stimulus (L) and on peak fluorescence analysis (L’, paired Student’s t test, shock-LDT: t(5) = 5.469, p = 0.0028, LDT: t(5) = 2.732, p = 0.0412); trial response detail depicted in heatmap (M). (N-O) Coca-LDT and LDT response to cocaine, showing an activity decrease aligned to cocaine injection, depicted in z-score of the DF/F on PSTH of the stimulus (N) but not on peak fluorescence analysis (N’), with trial response detail depicted in heatmap (O). (P-Q) Shock-LDT and LDT response to cocaine, showing activity decrease to cocaine injection, depicted in z-score of the DF/F on PSTH of the stimulus (P) but not on peak fluorescence analysis (P’); trial response detail depicted in heatmap (Q). Z-score data are represented as trial mean (darker line) ± SEM (lighter area). Lines over Z-score graphs indicate periods of significant difference from baseline as defined by bootstrap 95% CI. * p < 0.05, ** p < 0.01, *** p < 0.001

Exposure to TMA, a rewarding odor for the animals, induced a robust increase in activity across the LDT population as well as within both cocaine- and shock-recruited ensembles, as supported by the bootstrap 95% CI and peak fluorescence analysis (Figure 4B-E).

In contrast to this odor-evoked response, condensed milk consumption was associated with an overall reduction in activity across both ensembles and the LDT population overall (Figure 4F, H, G, I). Notably, peak fluorescence analysis revealed statistically significant differences between pre-stimulus and stimulus periods exclusively in the coca-LDT ensemble, indicating a more pronounced modulation in this population (Fig. 4F′, H′).

Unlike condensed milk, solid pellet consumption was associated with increase in activity across the LDT population and within both coca- and shock-recruited ensembles, also reflected in the peak fluorescence analysis (Figure 4J-M).

By contrast, cocaine administration produced only a minimal impact on the activity of these ensembles, with no statistically significant post-injection differences and no detectable stimulus-locked responses across populations (Fig. 4N–Q).

### Valence specific ensembles respond to conditioned place preference and fear conditioning

Our previous results showed that the LDT and its neuronal ensembles (LDT-coca and LDT-shock) respond differently to distinct stimuli. Previous studies have also shown that specific LDT populations or projections are involved in cocaine-induced conditioning^9,10^, and that activation of some LDT populations is sufficient to induce place preference ^2,3^. Based on this, we aimed to assess the responses of the LDT-coca and LDT-shock ensembles during cocaine conditioning.

To achieve this, we submitted mice to a cocaine conditioned place preference (Figure 5A). The analysis of time spent in each chamber showed that mice occupied the cocaine chamber longer after conditioning (Figure 5B-C). Hence, we analysed the GCaMP8m activity from both the coca-LDT and shock-LDT groups, and the LDT sRGCECO activity, extracting the number of wave peaks in 10 seconds bins (Figure 5D). This showed a peak frequency decrease in shock-LDT from pre-to post-conditioning in both cocaine and saline chambers (Figure 5E-F).

**Figure 5.**
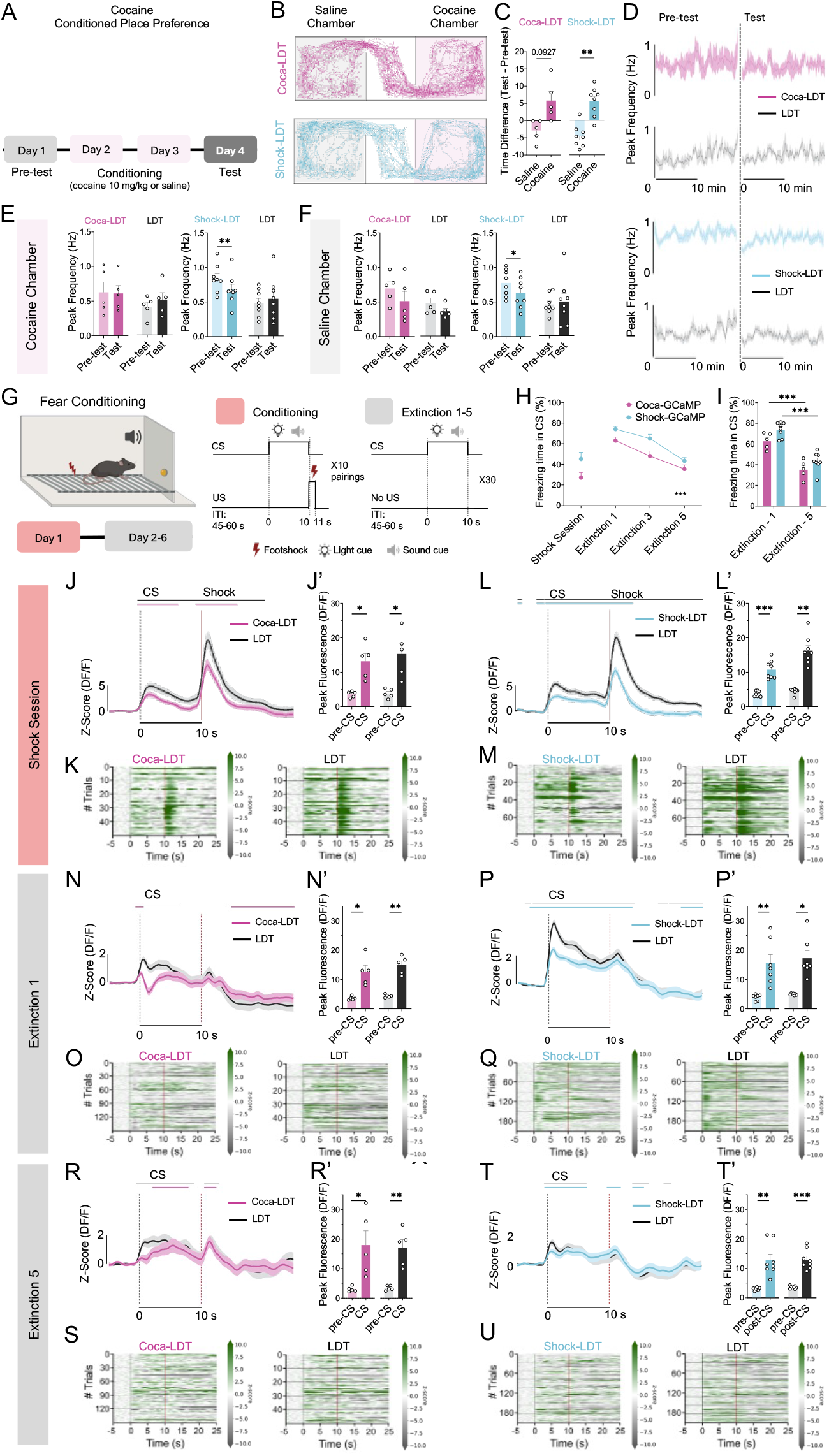
Cocaine and shock conditioning effect on LDT and shock- and cocaine-recruited LDT neuronal populations’ activity. (A) Graphical representation of cocaine conditioned place preference. (B) Example of mice movement tracings from coca-LDT (top) or shock-LDT (bottom) group across chambers on the test day. (C) Difference of time spent in cocaine and saline conditioned chambers on the test day in relation to the pre-test day (paired Student’s t test, coca: t(4) = 2.200, p = 0.0927, shock: t(5) = 3.555, p = 0.0163). (D) Example tracings of peak frequency (10 s bins) of DF/F activity of coca-LDT and LDT (top) or shock-LDT and LDT (bottom) along pre-test (left) and test (right) session. (E) Mean peak frequency on cocaine chamber during pre-test and test session, showing peak frequency decrease in shock-LDT from pre-test to test (paired Student’s t test, t(7) = 4.524, p = 0.0027). (F) Mean peak frequency on saline chamber during pre-test and test session, showing peak frequency decrease in shock-LDT from pre-test to test (paired Student’s t test, shock-LDT: t(7) = 2.713, p = 0.0301). (G) Graphical representation of fear conditioning test. (H-I) Freezing time during the CS on the several test days, showing decrease of freezing time during extinction days (two-way ANOVA, time factor: F(4, 44) = 15.20, p < 0.0001, *post hoc:* coca-LDT: Extinction 1 vs Extinction 3: t= 4.038, p = 0.0021, Extinction 1 vs Extinction 5: t = 6.909, p < 0.0001, Extinction 3 vs Extinction 5: t = 2.871, p = 0.0293, shock-LDT: Extinction 1 vs Extinction 5: t = 8.298, p < 0.0001, Extinction 3 vs Extinction 5: t = 5.827, p < 0.0001). (J-K) Coca-LDT and LDT response to CS and shock on the shock session, showing an activity increase aligned to the CS onset, depicted in both z-score of the DF/F on PSTH of the stimulus (J) and peak fluorescence analysis (J’, paired Student’s t test, coca-LDT: t(7) = 10.06, p < 0.0001, LDT: t(4) = 3.878, p = 0.0179), with trial response detail depicted in heatmap (K). (L-M) Shock-LDT and LDT response to CS and shock on the shock session, showing activity increase to CS onset, depicted in z-score of the DF/F on PSTH of the stimulus (L) but not on peak fluorescence analysis (L’); trial response detail depicted in heatmap (M). (N-O) Coca-LDT and LDT response to CS on the first day of extinction, showing an activity decrease aligned to CS, depicted in z-score of the DF/F on PSTH of the stimulus (N) but not on peak fluorescence analysis (N’), with trial response detail depicted in heatmap (O). (P-Q) Shock-LDT and LDT response to CS on the first day of extinction, showing activity decrease aligned to the CS, depicted in z-score of the DF/F on PSTH of the stimulus (P) and peak fluorescence analysis (P’, paired Student’s t test, shock-LDT: t(6) = 3.900, p = 0.0080, Wilcoxon test, LDT: W = 28.00, p = 0.0156); trial response detail depicted in heatmap (Q). (R-S) Coca-LDT and LDT response to CS on the fifth day of extinction, showing an activity decrease aligned to CS, depicted in z-score of the DF/F on PSTH of the stimulus (R) and on peak fluorescence analysis (R’, LDT: t(4) = 8.845, p = 0.0084), with trial response detail depicted in heatmap (S). (T-U) Shock-LDT and LDT response to CS on the fifth day of extinction, showing activity decrease aligned to the CS, depicted in z-score of the DF/F on PSTH of the stimulus (T) and peak fluorescence analysis (T’, Wilcoxon test, shock-LDT: W = 36.00, p = 0.0078, paired Student’s t test, LDT: t(7) = 8.347, p < 0.0001); trial response detail depicted in heatmap (U). Scalebars represent 200 µm. Z-score data are represented as trial mean (darker line) ± SEM (lighter area). Lines over Z-score graphs indicate periods of significant difference from baseline as defined by bootstrap 95% CI. Bar data are represented as animal mean ± SEM. * p < 0.05, ** p < 0.01, *** p < 0.001.

These data suggest LDT and its ensembles are not impacted by the cocaine conditioning, though the shock-LDT has a decreased frequency after conditioning to either cocaine or saline.

Regardless of this lack of significant impact in cocaine place preference, we still targeted the LDT-coca and LDT-shock response to a cue predicting an aversive event.

Mice performed a Fear conditioning task, consisting of a pure tone and light cue concomitant exposition for ten seconds (CS), right after which a 1 s footshock (US) was administered (Figure 5G) – conditioning sessions. The following five days were extinction sessions, with solely cues exposition. The freezing time during the CS decreased across the extinction days for both coca-LDT and shock-LDT (Figure 5H-I). As previously, both ensembles, as the LDT, responded to shocks, but also did they to the CS that predicted the footshock delivery, shown by the bootstrap 95% CI and the peak fluorescence analysis (Figure 5J-M). During the first day of extinction we registered also an activity increase during the CS exposition for both ensembles and the LDT, detected by the bootstrap 95% CI and the peak fluorescence analysis (Figure 5N-Q), a response that was still recorded on the fifth day of extinction (Figure 5R-U).

Overall, these results on the shock conditioning show that the LDT and both coca-LDT and shock-LDT ensembles were robustly activated by the footshock, and by the CS predicting shock, which was maintained during extinction.

### Modulation of cocaine-recruited LDT ensembles shifts preference

Considering the functional response of coca- and shock-recruited populations, we next aimed to understand whether optical re-activation of either ensemble is sufficient to change behavior. We injected TRAP2 mice bilaterally in the LDT with a cre-dependent AAV5 expressing activating opsin ChR2 (or YFP for control group) followed by fiber implantation in the same region (Figure 6A). Four weeks later, animals were TRAPped for either cocaine (leading to coca-ChR2 or coca-YFP groups) or shock (leading to shock-ChR2 or shock-YFP groups) stimuli (Figure 6A-B). Behavioural tests started 3 weeks after TRAPing to guarantee opsin expression (Figure 6B). Mice were first tested on a Real-Time Place Preference test (RTPP) to evaluate the reinforcing properties of specific optical stimulation of each ensemble (Figure 6C). Here, animals would receive optogenetic stimulation of the specific LDT ensemble every time and for the duration that the animal stayed on that side (chamber ON). Optical stimulation of cocaine- or shock-recruited LDT populations was not sufficient to significantly induce behavioral preference (Figure 6D-E).

**Figure 6.**
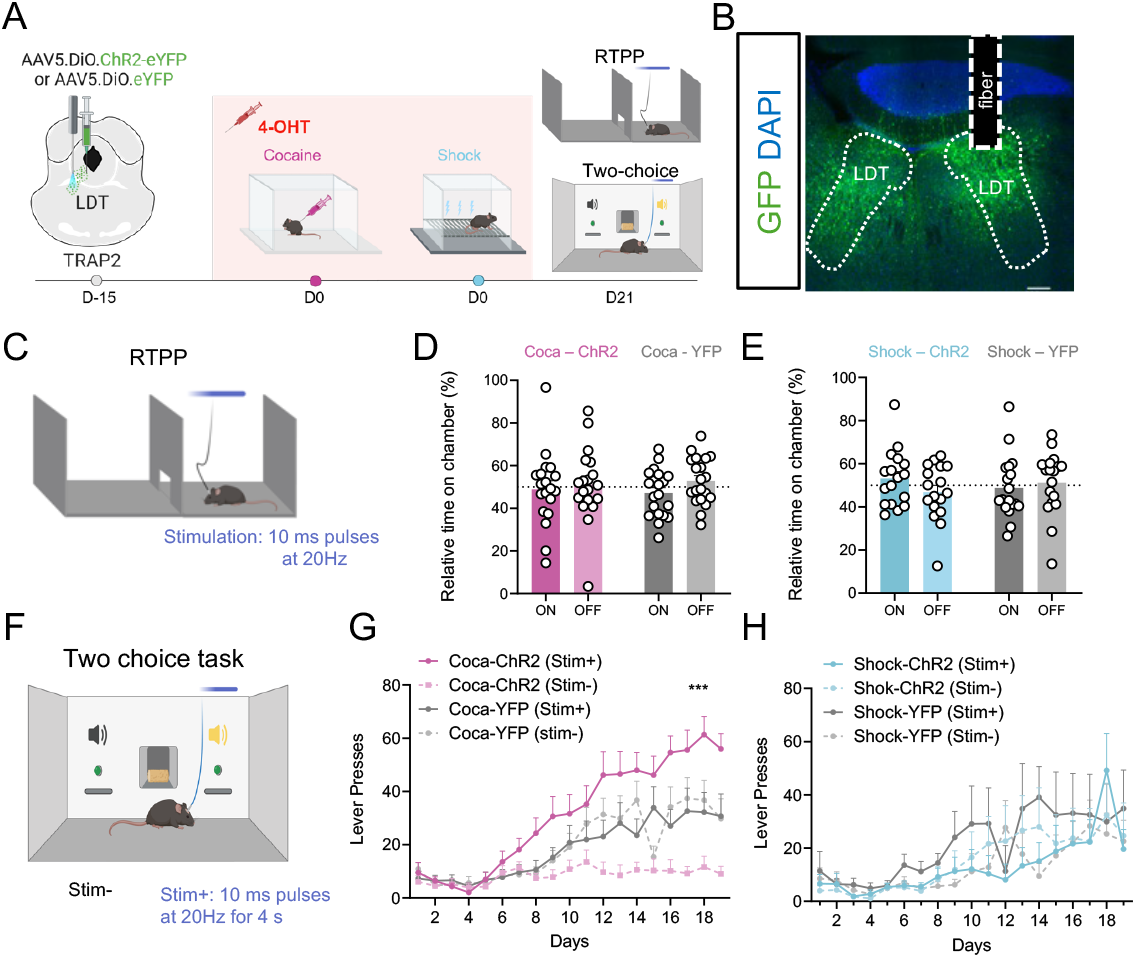
Modulation of coca-LDT and shock-LDT ensembles effect in preference. (A) Injection of cre-dependent ChR2 and YFP for control in the LDT of TRAP2 mice (n = 40) and strategy of recruitment of neuronal populations activated by cocaine (coca-ChR2: n = 10; coca-YFP: n = 10) or shock (shcok-ChR2: n = 10; shock-YFP: n = 10). (B) Histological checking of viral expression. (C) RTPP task representation, wherein mice were able to freely move between two similar chambers, the presence in one of them being coupled with light stimulation with blue light. (D-E) Preference analysis for the stimulated chamber for coca-ChR2 (D) or shock-ChR2 (E) neurons and respective YFP controls, showing no preference towards any of the chambers. (F) Two-choice operant task, where mice were presented to 2 sucrose pellet-giving levers, but pressing one of them having light-stimulation. (G) Presses on the stimulated (stim+) and unstimulated (stim-) levers for the coca-ChR2 (G) or shock-ChR2 (H) neurons and respective YFP controls, showing a preference towards the stim+ lever on coca-ChR2 mice (two-way ANOVA, group effect: F(1.907, 86.88) = 14.10, p < 0.0001). (J) Presses on stim+ and stim-levers for the shock-recruited LDT population and controls, showing no diferences in preference. Scalebars represent 200 µm. Data is presented as mean ± SEM. * p < 0.05, ** p < 0.01, *** p < 0.001

To assess whether each ensemble reactivation was sufficient to modulate reward-related behaviour, we tested mice on an operant two-choice test under a fixed ratio of 1 (FR1). Briefly, mice are exposed to two active levers, both delivering one food pellet reward and one of the levers also triggered a 4s blue light stimulation (Stim+; excitation: 10 ms pulses, 10 Hz) (Figure 6F).

Mice receiving optical activation of cocaine ensemble (coca-ChR2) showed a clear preference for stim+ lever in comparison with the stim-lever (Fig. 3K) across acquisition days, significantly exceeding YFP group (Figure 6G). No preference for either lever was observed in the coca-YFP group (Fig. 3K-L). Interestingly, optical activation of shock-ensembles did not induce any significant differences between lever in the shock-ChR2 nor between the shock-YFP group (Figure 6H).

Together, these results indicate that activation of a cocaine ensemble in the LDT can reproduce reward-associated behavioral responses, supporting their role in driving preference and reinforcement value.

## Discussion

Though little is still clear about the LDT role in valence encoding, it has been stated repeatedly that it or some of its projections activation leads to preference shift in preference-assessing tasks ^2–4,7,12,16^. This seems to be in line with our results, as we found that sequential exposure twice to the same stimulus led to a higher colocalization of activated neurons on the first and second exposure comparing to when animals were exposed to stimuli of different valence. Single exposures to cocaine or footshock label LDT ensembles with limited cellular overlap, supporting the presence of stimulus-defined representations within this brainstem nucleus. Both ensembles were neurochemically heterogeneous, with a predominance of GABAergic neurons, consistent with prior work indicating functional diversity within LDT inhibitory circuits. For instance, a study on GABAergic neurons in the LDT has described that a group of these may be activated by aversive stimuli, whereas others are responsive to rewarding stimuli (Du et al., 2023).

Optogenetic activation of LDT cholinergic and glutamatergic projections to the NAc increased preference in different testing paradigms ^3^ and the overall activation of LDT cholinergic cells also led to preference instatement ^17^. Another study found that optogenetic inhibition of LDT glutamatergic neurons leads to reduced cocaine-induced locomotor activity ^18^. Nonetheless, and in contrary to what had been described previously, a study suggested that the cholinergic LDT to thalamus projection optogenetic activation during an anxiety test (elevated plus maze) increased anxiety-like behaviour ^16^, opposing the common findings on the role of the LDT cholinergic populations in reward mediation. The GABA population was perhaps the upmost studied LDT ensemble regarding differences in valence response and led to the most promising results. For instance, inhibition of GAD2 LDT-VTA projection terminals reduced nicotine-led aversion ^19^. In addition, activation of both PV^+^ and SOM^+^ GABA interneurons in the LDT reduced responses to a fear-conditioning protocol ^6^. This is consistent with our findings that specific LDT cell activation can modulate preference. Regarding the differences of projection targets of GABA LDT cells, previous studies from our team showed that the activation of GABA LDT-NAc projections shifted preference to the non-stimulated lever in a two-choice operant task ^3^. This is similar to what has been found in the LDT-VTA GABA projections, for chemogenetic excitation decreased reinstatement of cocaine self-administration ^20^ and increased freezing response to footshocks ^21^. For the contrary, chemogenetic inhibition reduced freezing in response to footshocks ^21^. These dichotomies within each population responses aligns with our findings, for we found that the same shock-recruited LDT neuronal ensembles were activated by physical aversive events and by an aversive odour, which may even underlie the whole LDT population activation.

Cocaine-recruited ensembles responded to the physical or olfactory events. Although ensemble recruitment was biased by the TRAPing stimulus, ensemble responses during subsequent testing were not strictly valence selective. Both cocaine- and shock-tagged ensembles were robustly engaged by highly salient aversive stimulation and by conditioned threat cues, and both responded to odor stimuli, indicating that LDT ensembles can be broadly sensitive to salience and sensory modality. This pattern suggests that TRAPing captures ensembles with a history of recruitment by a particular experience, rather than immutable “valence-labeled” units. A key implication is that ensemble identity may reflect convergent inputs engaged during the tagging event and the state of the animal, which can generalize across stimuli sharing arousal or motivational relevance.

Actually, evidence from other brain structures also supports the concept of neuronal responses, which depend on the input information that controls the structure’s activity. Notably, the Amygdala contains distinct neuronal populations that respond to positive and negative stimuli, which also acquire predictive responses during learning^22^. Similarly, the VTA, that the LDT projects to, has been found to have distinct populations mediating reward and aversion depending on their inputs – those receiving projections from the LDT typically mediate reward, while those receiving inputs from the PFC mediate aversion^23^. Thence, the segregation of valence-encoding populations in the LDT mirrors broader brain regions involved in reward and aversion processing. Still, further research on the field is necessary to clarify the distinct function of LTD populations role in specific valence encoding.

Despite shared recruitment by salient events, reactivation of cocaine-tagged ensembles biased action selection in an operant choice task, whereas reactivation of shock-tagged ensembles did not produce avoidance or place preference. This is quite new on the field, as most studies focus on the neurotransmitter content to distinguish populations and not which stimuli are capable of activating them. As previously described, it was known so far that cholinergic and glutamatergic populations chiefly could induce preference ^3,12,16,17,24^, and GABA LDT populations were more evidently diverse on the kind of response their activation could elicit ^6,7^. Activating the recruited population we could better describe the effect of the interplay of the different ensembles present within each one based on the neurotransmitter content, leading, broadly, to a preference encoding in an operant task, whereas the interplay of all the neurotransmitter based populations recruited by the shock event on the moment of TRAPping did not affect, in general, preference in the same kind of preference test. One possibility is that cocaine-tagged ensembles preferentially engage downstream circuits that support instrumental invigoration or action reinforcement, whereas shock-tagged ensembles may encode salience without being sufficient to drive avoidance in the absence of threat contingency. The dissociation between the lack of real-time place preference and the presence of operant bias further suggests that ensemble activation interacts with task structure, potentially by reinforcing action–outcome associations rather than generating a simple hedonic state.

Together, our findings support a model in which the LDT contains experience-defined ensembles that are recruited by salient events and can influence motivated behavior in a context-dependent manner. Defining the projection targets and synaptic mechanisms through which cocaine-tagged ensembles bias choice, and determining how ensemble recruitment evolves with learning, stress or repeated drug exposure, will be important directions for understanding how brainstem circuits contribute to maladaptive decision-making.

## Methods

### Animals and Treatments

Male and female homozygous ‘Targeted Recombination in Active Populations 2’ (TRAP2) mice (Fos^2A-iCreER/2A-iCreER^/J; stock #030323, Jackson Labs) were used for optogenetic and photometry studies. This model expresses Cre recombinase under the c-fos promotor, allowing a cre-dependent virus to be expressed in stimuli-activated cells under the presence of 4-hydroxytamoxifen (4-OHT) ^25,26^.

Animals were kept under standard laboratory conditions (light/dark cycle of 12h/12h; 22ºC), food and water available *ad libitum*, unless described otherwise. Behaviour experiments were undergone on the light period. All protocols involving mice were approved by the Ethics Committee of the Life and Health Sciences Research Institute (ICVS) and by the national competent entity, the Direcção-Geral de Alimentação e Veterinária (DGAV) (#8332, dated of 2021-05-08). Every technique was performed following established ethical and legal standards, including the European Union Directive 2010/63/EU. Health monitoring followed FELASA guidelines, thus confirming the Specific Pathogen Free health status of sentinel animals housed in the same room. All experimenters, animal facilities’ structures and caretakers were accredited by DGAV. Every possible effort was made to minimize the number of animals used, and their discomfort and suffering.

### Stereotaxic surgeries and fibre implanting

TRAP2 mice used for optogenetics and photometry were anaesthetised until verified lost paw withdrawal reflex with 8% sevoflurane (SevoFlo, Chicago, IL, USA), in oxygen, at a rate of 0.8-1 L/min. During surgical procedures anaesthesia was provided on a nosecone (2-3%; 0.8-1 L/min), maintaining body temperature at 37ºC using a heating pad. Pre- and post-operative (6h after surgery, and afterwards, daily up to 3 days, if animals showed signs of discomfort) analgesia was carried with buprenorphine (i.p.; 0.05 mg/kg; Bupaq, RichterPharma, Austria). Signs of dehydration or reduced movements were handled with 0.9% saline (i.p.; 20 μL/g) or a multivitamin supplement (i.p.; 1 : 10 in saline 0.9%; 20 μL/g; Duphalyte, Zoetis, Spain), respectively, daily, up to 3 consecutive days.

Animals used in photometry were injected with 400 nL of a mix 1:1 of cre-dependent AAV5-syn-Flex-GCaMP8m and general expressing AAV2/5-CAG-sRGECO in the LDT (from Bregma^27^: -5.2 mm anteroposterior (AP), ±0.6 mm mediolateral (ML) and -3.4 mm dorsoventral (DV)). For optogenetic experiments mice were injected bilaterally (300 nL each side;1 nL/s) with AAV5-DIO-CHR2-eYFP or AAV5–EF1a–YFP for control in the LDT (from Bregma ^27^: -5.2 mm AP, ±0.6 mm ML and -3.4 mm DV). Injections were made using a Nanoject III injector (Drumond Scientific Company, Broomall, PA, USA), with the micropipette being immobile for 5 mins after injection to allow viral diffusion. After viral infusion, animals were implanted with fibres using the same coordinates ^27^, but with -3.3 mm DV. Implants had Ø 1.25 mm ferrule with a multimode optic fibre with core of Ø200 µm and 0.22 Numerical Aperture (NA; for optogenetics using a set from ThorLabs, Newton, NJ, USA; for photometry, a Doric Lenses set, Québec, Canada). Ferrules were secured to the skull with dental cement (C&B kit, Sun Medical, Shiga, Japan).

### TRAP Induction

For TRAP induction, 3 – 4-month-old mice were habituated for 7 days to the switched-off operant apparatus (mice to be induced with shock or exposed to sucrose), saline 0.9% i.p. injections and then to single-housing for 6h. The next day, mice were exposed to cocaine hydrochloride (20 mg/kg, i.p.; 2mg/mL in saline 0.9%; # C5576-1G, Sigma-Aldrich, St. Louis, MO, USA); saline 0.9% (i.p.); a shock session (1.5 mA every 30 seconds, total 20 footshocks, 10 min sessions; house light on for the whole session) in an operant chamber with a shocker module (Med Associates, St. Albans, VT, USA); to the turned off shock apparatus (house light on); the operant chamber delivering sucrose 20% or water for control (50 µL, 180 solution deliveries, 90 min session). Right after stimulation, animals were injected with 4-OHT (50 mg/kg; #H6278, Sigma-Aldrich, St. Louis, MO, USA) and single-housed for 6 hours to avoid overflow of social stimuli. The 4-OHT solution was prepared by dissolving it in 100% ethanol (52ºC, 15-20 mins under shaking). Castor oil and sunflower seed oil mixture (1:4) considering the 4-OHT final concentration of 5 mg/mL, at room temperature (RT). This solution was prepared on the day it was used and kept at 36ºC before administration.

### Endogenous c-Fos labelling

On the experimental day, following the habituation period similar to TRAP induction, the animals went through the respective stimulus session and then isolated in individual cages (after 45 minutes OF session in the Saline and Coca groups), being sacrificed approximately 90 min after to detect endogenous c-Fos protein induced by the session. For the sucrose group, the peak of consumption was considered in the middle of the session therefore, the animals were sacrificed 45 min after exiting the apparatus.

### Behavioural Experiments Odours Exposure

Animals were habituated to an empty white opaque plastic arena for 3 days (53 cm x 35 cm x 20 cm, base similar to home cages bedding), underneath a chemical hotte (Ultragene, Santa Comba Dão, Portugal) and connected to the photometry system. Animals were then exposed to trimethylamine (TMA; 10mM, diluted in water) or trimethylthiazoline (TMT; 1.2 mM, diluted in water). Odour exposure was counter-balanced between groups within and across each of two days of exposure. Exposures consisted on a cotton swab being immersed in the odour solution and then approached to the mouse nose as close as possible ensuring it would not touch (15 trials; random inter-trial intervals (ITI) of 45-70 s). Photometry recordings started 10 mins before expositions to stabilise signal. The whole procedure was done underneath a portable chemical hotte (Ultragene, Santa Comba Dão, Portugal) to ensure the extraction of remaining smell in the arena.

### Tail Pinch and Tail Lift

On the same arena used in odours exposure, mice were pinched with tweezers and the next day lifted by the experimenter, for 15 times, with a random ITI of 45-70 seconds. The photometry recordings started 10 minutes before expositions so signal could stabilise.

### Liquids Exposure

Animals were habituated to the operant chamber and the patch cable. The box (17.8 cm × 19 cm × 23 cm; metal floor with a central magazine disposing liquids delivered by a solenoid) was placed within a sound-attenuating box and controlled by pyControl software. House light was kept on during every session. After 2 days of chamber habituation, photometry recordings started during which mice were exposed to a different liquid every day (60 exposures, 15-35 s ITI), on the following order: water (mice were water-restricted for 12h before exposure), sucrose 10% wt/vol in water, sucrose 30% wt/vol in water, quinine 1mM (Sigma-Aldrich, St. Louis, MO, USA) diluted in water (after a day interval with exposure to sucrose 10% wt/vol in water). During the whole liquid exposure protocol, mice were food restricted.

### Real-time Place Preference (RTPP)

RTPP apparatus consisted on a 60 cm x 50 cm x 40 cm plexiglass arena divided in half. The resulting two chambers were connected by a central opening. The animals were habituated to the chambers connected to the optogenetics cable for 20 mins and tested on the following day. Each animal was placed in the centre opening and left to freely move between the two chambers for 20 mins. While the animal was in one of those chambers, optogenetic stimulation was activated manually and continuously (chamber ON – randomly assigned to each animal and balanced across groups; light stimulation of 10 ms pulses at 20 Hz). Animals’ behaviour was recorded with a camera, and the time spent in each chamber was registered. Analysis consisted on the relative time spent in chamber ON considering the whole test time.

### Conditioned Place Preference (CPP)

Mice were habituated with the patch cable connected for 2 days to the CPP apparatus (50.5 x 16 cm), consisting of two conditioning chambers (20 x 16 cm), separated by a middle neutral one (10.5 x 16 cm). During the pre-test and test (1 day each, on 20 min session), animals could freely move between chambers, and the time spent in each conditioning chamber was registered. In between, mice were submitted to 2 conditioning days, in which animals were injected with 0.9% saline and confined to one of the conditioning chambers and injected with cocaine (10mg /kg, i.p., #C5576-1G, Sigma-Aldrich, St. Louis, MO, USA) and confined to the other conditioning chamber for 30 min. Every animal was conditioned once with saline and cocaine on the two days and these conditioning were separated by 5 hours minimum; conditioning chamber attributed to saline or cocaine was counterbalanced across groups.

The animal locomotion was recorded during all the sessions. Photometry signals were recording during the pre-test and test sessions, and during cocaine conditioning of the first day and saline conditioning of the second day, or vice-versa, counterbalanced across groups.

### Fear Conditioning

Mice were habituated to the were familiarized to the behaviour apparatus (17.8 cm × 19 cm × 23 cm; metal grid floor) for 2 days with the patch cable connected, and houselight on. Next, mice performed a shock conditioning session, which started with 60 s period habituation with only the houselight on. They were then exposed to CS (80-dB, 2 kHz tone and a cue light) after which the US consisting of a mild foot shock (0.5 mA, over 1 s) was administered through the grid floor. Trials were separated by random ITI (45–50 s), on a total of 10 trials. The next five days consisted on extinction sessions, in which mice performed 30 trials with only CS exposure, but without US delivery. All sessions were recorded in video and photometry recordings were acquired. We measured freezing during the CS period from all sessions by video observation by an experimenter blind to group allocation, consisting on the time mice spent immobile and calculated as percentage of total cue time (freezing time/total cue time x 100).

### Two-choice Reinforcement

Animals were habituated for two days to the operant boxes with only house light on (21.59 cm x 18.08 cm x 12.7 cm; Med Associates, St. Albans, VT, USA).

During reinforcement schedule, food-restricted mice were trained to press two levers that dispose a sucrose food pellet on a central portico (levers on opposite sides of the central portico, on a fixed ratio 1 (FR1)). Each lever had a 4s sound cue associated (tone or white noise) and was illuminated. Pressing one of the levers led to optogenetic stimulation (80 light 10 ms pulses delivered at 20 Hz over 4s) – stim+ lever. The assignment of the stim+ lever was balanced between sound cues associated and across groups. The first two trials had the levers presented one at a time, after which both levers were available simultaneously. The total number of lever presses in each one of the lever types was registered for a total session time of 20 mins.

### Fibre Photometry Recordings

The neuronal ensembles activity (cocaine or shock recruited or general LDT populations) was recorded using a Doric Lenses acquisition system (Doric Studio, Quebec, Canada). An optic fibre patch cable (Ø200 µm core and 0.22 NA 1 m wide; Doric Lenses, Quebec, Canada) was coupled to the implanted optic fibre with an ADAL3 interconnector for Ø1.25 mm ferrules (ThorLabs, Newton, NJ, USA). The system used three LEDs (CLED_465; CLED_405; Doric Lenses, Quebec, Canada) via a LED driver. Excitation light with three different wavelengths of 465 nm for GCaMP6f, 405 nm for the GCaMP6f isosbestic channel, and 561 nm for jRGECO1a, and an emitted light was passed through a minicube [ilFMC6-G2_IE(400–410)_E1(460– 490)_F1(500–540)_E2(550–580)_F2(600–680)_S; Doric Lenses, Quebec, Canada]. GCaMP6f and jRGECO1a signals and a movement control signal from the GCaMP6f isosbestic channel were recorded using Doric Studio Software through a console. A more directed control to the jRGECO1a signal, an isosbestic, was not available. The sensors light emitted is transmitted through the same optic fibre, split from the illumination light by optical filters and converted to electrical signals by a high sensitivity photoreceiver (Newport 2151 with lensed FC adapter; Doric Lenses, Quebec, Canada). For liquids exposure, the digital inputs were read at the same sampling rate as the analogue signals from the behaviour box to enable post hoc synchronization of the photometry data with the behavioural hardware. The system uses a digital input to receive a time-to-live synchronisation (TTL) signal each time a reward was made available. For odours’ exposure, tail pinch and lift, the TTL signal was manually marked on each trial event.

### Fibre Photometry Data Analysis

Photometry data was initially performed using a free open-source package ‘Guided Photometry Analysis in Python’ (GuPPy; #SCR_022353; ^28^). GuPPy used the control channel, the isosbestic channel, which includes primarily small movement artifacts and photobleaching, and subtracted from the signal. This resulting signal was used to normalise data from the baseline fluorescence, also dividing it by the control channel, then getting ΔF/F. Posteriorly, post-stimulus histograms (PSTH) were calculated on a defined window around each event: -10s to 15s from odour exposures, tail pinch and lift and -3s to 8s from liquid consumption (a poke and at least 2 lick events registered by the behavioural box within less than 250 ms after liquid delivery). We then used Python packages (Python version 3.9.12, Numpy version 1.21.5; #SCR_006903) to calculate the *z* score from the data present on the PSTH, by the difference of the mean value of baseline activity (considered from the minimum limit of event window to 0 s timepoint) for each trial of each animal and dividing by the standard deviation of the baseline activity. A bootstrapping confidence interval procedure was used with 95% CI to determine changes fluorescence changes within the PSTH timeframe, then expanded by a factor of √(*n*/(*n* − 1)) to account for narrowness bias, and only periods that were continuously significant (greater or less than the expanded BtsCI limits) for at least 0.5s were identified as significant ^29,30^. The PSTH’s area under the curve (AUC; using Python package Scikit-learn version 1.1.1 [sklearn. metrics. auc]) in specific periods of time were then calculated. The intervals used for data were [-2, 0]s and [0, 2]s. The whole data analysis procedure was done for both the green GCaMP6f and red jRGECO1a signals.

### Histology

### Euthanasia and Brain Sectioning

After all procedures, animals were deeply anaesthetised with a mixture of ketamine and medetomidine (respectively 150 mg/kg and 0.3 mg/kg, i.p.) and transcardially perfused with 0.9% saline, followed by a 4% paraformaldehyde (PFA) solution (pH 7.4 in Phosphate-buffered saline (PBS)). Brains were removed and post-fixed in PFA for 24h (mice used in optogenetics), or removed after whole heads immersion for 48 h in 4% PFA with fibre implant (mice used in photometry). After post-fixation, brains were immersed in sucrose 30% at 4ºC until being sectioned.

Sectioning was done on a vibrating microtome (VT1000S, Leica Biosystems, Nussloch, Germany), coronally at 40 μm thickness and slices stored in PBS at 4ºC. Sections of interest were chosen according to the *Mouse Brain Atlas* ^27^. For long-term storage, sections were cryopreserved with a cryoprotectant solution and kept at -20ºC.

### Immunostaining

Sections were washed 3 times for PBS with Triton-X100 (0.3%) (PBS-T) to permeabilise membranes and then blocked with 10% foetal bovine serum (FBS; in PBS-T at RT; Invitrogen, Waltham, MA, USA), to avoid unspecific binding. The primary antibody (AB) was incubated overnight at 4ºC. For sections of the fibre photometry experiment, there were two distinct molecular targets, so primary AB incubations were sequential. The following day, sections were washed with PBS-T (3x, 10 mins each) and the secondary AB was added, being incubated at RT for 2h. All antibodies were diluted in PBS-T containing 2% FBS. To identify cell nuclei, sections were washed twice in PBS-T and once in PBS, then being incubated with 4’,6-diamidino-2-phenylindole (DAPI; 1:1000, 10 mins, RT; Invitrogen, Waltham, MA, USA). At the end, they were mounted on glass slides (SUPERFROST PLUS, Thermo Scientific, Waltham, MA, USA) using Permafluor (Invitrogen, Waltham, MA, USA). Slides were stored under protection from light, at 4ºC. Primary antibodies consisted on rabbit anti-GFP (1:750, Ab290, Abcam, Cambridge, UK), goat anti-TdTomato (1:750, HBT020-2201, HenBiotech, Coimbra, Portugal). Secondaries used were Donkey anti-rabbit (1:100,Alexa Fluor 488, Invitrogen, Waltham, MA, USA) and donkey anti-goat (1:1000, Alexa Fluor 594, Invitrogen, Waltham, MA, USA).

### Image Acquisition and Analysis

To assess the localisation of virus injection and fibres placement, images from the LDT region were collected in an inverted fluorescence microscope (Olympus widefield inverted microscope IX81, Tokyo, Japan) on 10-20x objective magnification. About 6-10 slices per animal were used for analysis. On the slices where the fibre implant were detected, as well as virus injection, the coordinates were classified according to the *Mouse Brain Atlas* ^27^.

### Statistical Analysis

Statistical Analysis was performed in GraphPad Prism v10.3.0 (#SCR_002798, La Jolla, CA, USA), unless stated otherwise. Outlier detection was performed using Tukey’s Fences for Outliers test. The normality was checked using Kolmogorov-Smirnov test (KS), and parametric tests used if KS>0.05. Comparison between two groups was done using unpaired or paired Student’s t test, properly signed; or Mann-Whitney (unpaired) or Wilcoxon (paired) tests whenever data normality was not achieved. Comparison of three or more groups was done using one-way analysis of variances (ANOVA), followed by Bonferroni’s *post hoc* multiple comparison test to determine group differences. Two-way ANOVA was used to compare groups with more than one measurement, followed by Bonferroni’s *post hoc* analysis for multiple comparisons. Results are presented as mean ± standard error of the mean (SEM). Statistical significance was accepted for p < 0.05.

---

**Supplementary Table 1.**
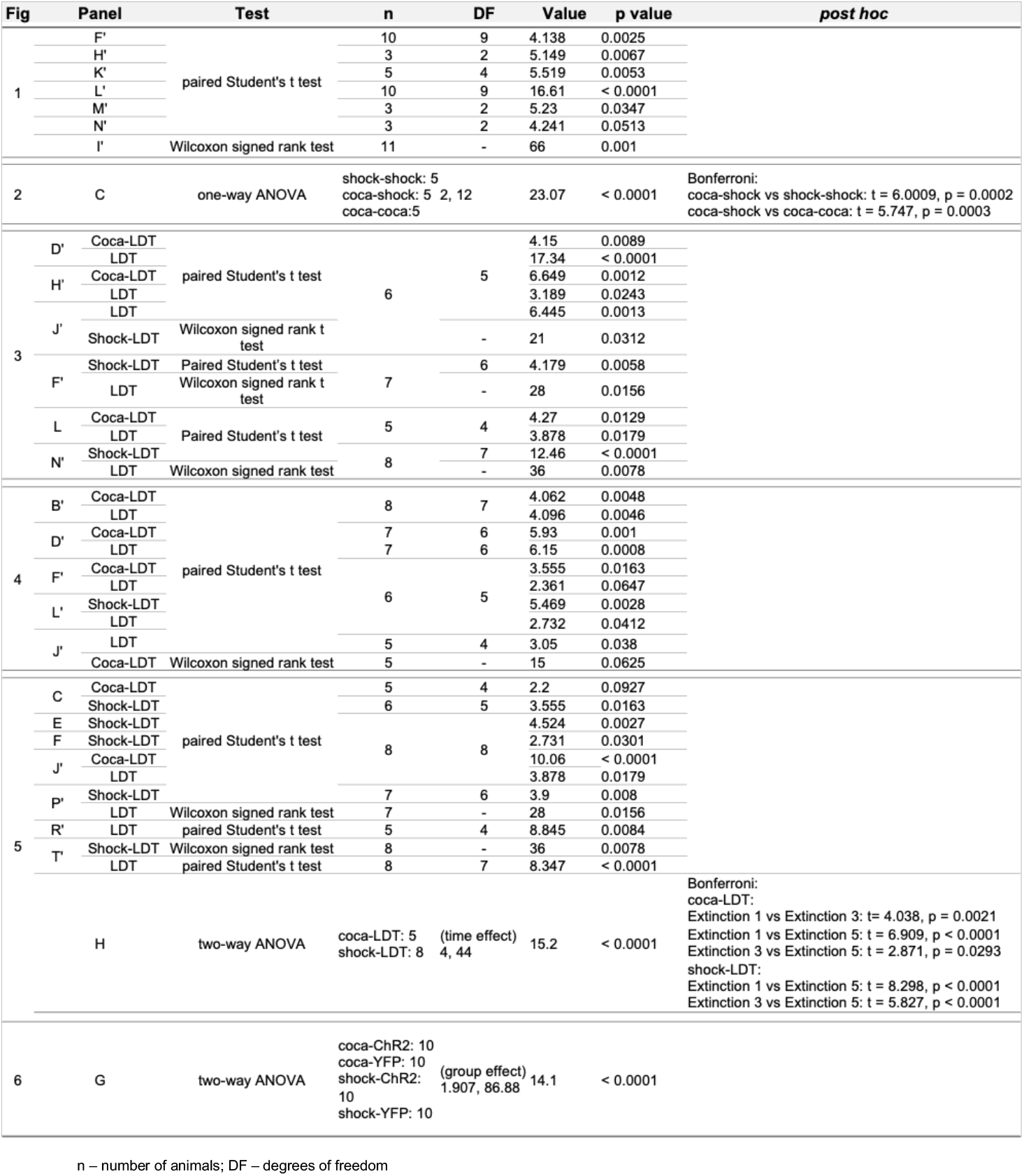
Statistic analysis and results used along the study.

